# Antibody binding epitope Mapping (AbMap) of two hundred antibodies in a single run

**DOI:** 10.1101/739342

**Authors:** Huan Qi, Mingliang Ma, Chuansheng Hu, Zhao-wei Xu, Fan-lin Wu, Nan Wang, Dan-yun Lai, Yang Li, Shu-juan Guo, Xiaodong Zhao, Hua Li, Sheng-ce Tao

## Abstract

Epitope mapping is essential for the understanding how an antibody works. Given millions of antibodies short of epitope information, there is an urgent need for high-throughput epitope mapping. Here we combined a commercial phage displayed random peptide library of 10^9^ diversity with next generation sequencing to develop <u>A</u>ntibody <u>b</u>inding epitope <u>Map</u>ping (AbMap) technology. Over two hundred antibodies were analyzed in a single test and epitopes were determined for >50% of them. Strikingly, the antibodies were able to recognize different proteins from multiple species with similar epitopes. We successfully identified the epitopes of 14 anti-PD-1 antibodies, including Sintilimab (*i.e.*, L128, A129 and P130), and confirmed that the binding epitopes of Nivolumab and Sintilimab are very close to the binding interface of PD-1 and PD-L1. The predicted conformational epitopes of Pembrolizumab and Nivolumab are consistent with their antibody-antigen co-crystal structures. AbMap is the first technology enables high-throughput epitope mapping.

**Highlights:**

- The first technology enables epitope mapping of two hundred antibodies in a single run
- Linear epitope was determined for >50% of the antibodies
- Distinct epitopes of 14 anti-PD-1 antibodies, including Sintilimab, were determined
- The predicted conformational epitopes of Pembrolizumab and Nivolumab are consistent with the known antibody-antigen co-crystal structures

---

Antibodies are a group of naturally existing proteins, which could specifically bind to antigens, especially proteins. Taking advantage of the high specificity of antibody-antigen binding, antibodies are widely applied as reagents for many different assays, *e.g.*, Western blotting, ELISA, flow cytometry and immunoprecipitation. According to CiteAb^1^, there are 4,494,439 antibodies by August 17, 2019 from 190 suppliers (https://www.citeab.com/). This number does not include the large number of home-made antibodies by researchers around the world. In addition, the market of antibody is growing rapidly, with an annual rate of 6.1% to 2025 (https://www.grandviewreresearch.com). From the first therapeutic antibody, *i.e.*, Muromonab OKT3, we have witnessed the sharp growth of this market. There are 70 FDA and EMA approved therapeutic antibodies on the market by 2017^2, 3^, for the treatment of a variety of diseases, from cancer, autoimmune disease^4^ to pathogenic bacteria^5^. In addition, there are over 570 therapeutic antibodies at various clinical phases, including 62 in late-stage^3^.

The binding of an antibody to an antigen is through epitope, which is a small area or a few amino acids on the antigen. Typically, there are two types of epitopes, *i.e.,* linear epitope and conformational epitope^6^. Epitope is one of the most critical parameters of antibody and antigen binding. By revealing the epitope, when using an antibody as reagent, it will make the experimental conclusions clear and relevant. For a therapeutic antibody, by knowing the epitope, it will help us to understand the precise mechanism of action. More importantly, it will facilitate the developer to protect the intellectual property at the right time.

There are several technologies for epitope mapping, generally, these technologies could be categorized into two types, *i.e.*, structure-based and peptide-based^7^. Structure-based technologies are the golden standards, including X-ray crystallography^8^, Nuclear Magnetic Resonance^9^, and more recently, Cryo-EM^10^. By August 5, 2019, there are 840 antibody-antigen co-crystal structures in PDB (www.rcsb.org). For any antibody, based on the antibody-antigen co-crystal structure, comprehensive binding interface is readily to be obtained. However, it is difficult to acquire high-quality antibody-antigen co-crystal, which is the key step of successful X-ray crystallography. It is even harder when the antigen is a membrane protein^11^. Peptide-based technologies include peptide microarray^12^, SPOT^13^, and phage or *E.coli* displayed peptides or protein fragments^14^. Typically, peptides cover a single or a set of antigens were synthesized and immobilized on a planar microarray, or present on phage/*E. coli*. The peptides that an antibody binds to are then enriched and identified. Usually, large number of peptides are required, and the construction of the microarray and the library is sophisticated. Compared with conformational epitope, linear epitope is much easier to be obtained by peptide-based technologies. In addition to the structure-based and peptide-based technologies, other technologies, including site specific mutagenesis^15^, and mass spectrometry^16^, and hydrogen-deuterium exchange mass spectrometry (HDX-MS)^17^, are developed. All these technologies are time-consuming, labor intensive and costly. The epitopes could only be resolved in a case by case manner. The success of one pair of antibody-antigen is difficult to be repeated on another pair. Only one or a set of antibodies could be analyzed a time, and none of them are suitable for high-throughput analysis.

With millions of antibodies are on the market, and epitope information are missing for most of them, we are aiming to develop an enabling technology to massively map the epitope of antibodies. Once the epitopes are mapped, lots of possibility could be explored. The schematic of our strategy is depicted in Figure 1, and we name it as AbMap. Key components of AbMap are phage displayed random peptide library, and the power of next generation sequencing (NGS) for decoding barcoded and mixed samples in an unprecedented way. Our underlying hypothesis is that the highly diversified phage displayed random peptide library could cover many, if not all, of the peptides that could be recognized by any given antibody.

**Figure 1.**
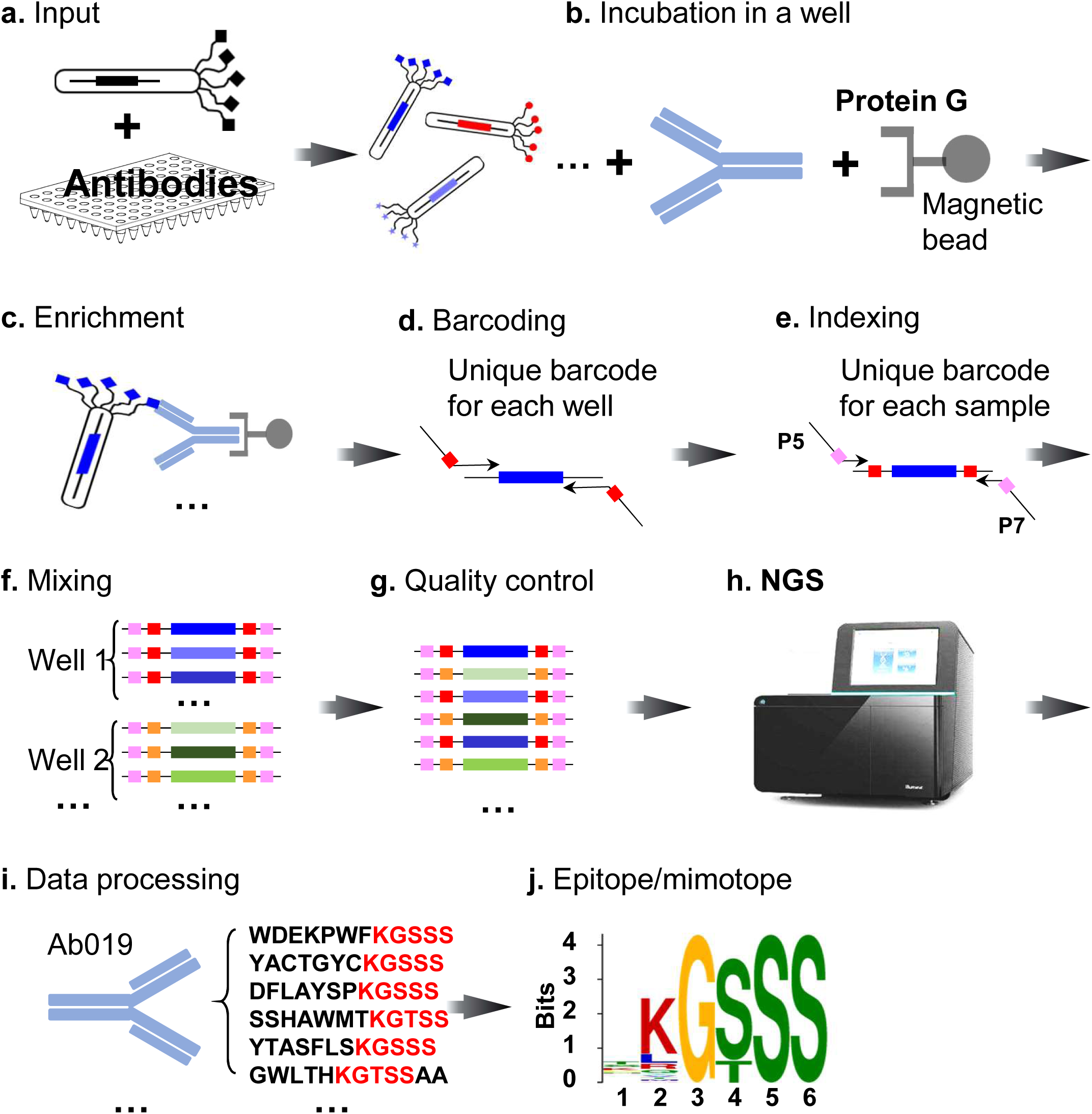
The schematic and workflow of AbMap. **a**. Antibodies are incubated with phage displayed random peptide library. **b-c**. The phages bound by the antibody are enriched by Protein G coated magnetic beads. **d**. The DNA is released and barcoded with unique barcode-pair in each well. **e**. A unique index is selected for each sample. **f.** The PCR products are mixed. **g**. Quality control of the barcoded and indexed PCR products. **h**. NGS analysis of the samples. **i**. Data processing. **j**. Epitope/mimotope calculation.

Taking several affinity-tag specific antibodies as examples, we first proved the feasibility of AbMap. The high-throughput protocol was then established and the capability was demonstrated by testing 202 antibodies in a single run. The results showed that linear epitopes could be determined for >55% of the antibodies. The identified peptides/epitopes were then explored for other interesting applications. Finally, a set of anti-PD-1 antibodies, including Nivolumab^18^, Pembrolizumab^19^ and Sintilimab^20^ were analyzed by AbMap. For different antibodies, distinct binding epitopes on PD-1 were determined, which are consistent to the conformational epitopes determined by X-ray crystallography.

## Results

### Schematic diagram of AbMap

The schematic and workflow of AbMap is shown in Figure 1. Antibodies (monoclonal or polyclonal) are aliquoted in 96-well plates (Figure 1a), phage displayed random peptide library, in our case, Ph.D-12, are incubated with the antibody in each well (Figure 1b). The phages bound by the antibody are enriched by Protein G coated magnetic beads (Figure 1b and Figure 1c). The DNA is released from the enriched phages, and PCR product is barcoded with unique barcode-pair in each well (Figure 1d), and a unique index is chosen for each batch of antibodies (Figure 1e). The barcodes and indexes used in this study were shown in **Table S1**. To avoid the possible amplification bias, the index is incorporated for each antibody individually in each well, and then mixed (Figure 1f). To control the sample quality, *e.g.*, the purity and the concentration of the DNA samples, and the integrity of the PCR products are measured (Figure 1g). The sample is then subjected to NGS (Figure 1h). The sequencing data is processed through filtering, normalization and *etc* (Figure 2). The peptides that are strongly bound by each antibody are determined (Figure 1i). Epitope or mimotope of a given antibody is revealed by a variety of computational tools (Figure 1j).

**Figure 2.**
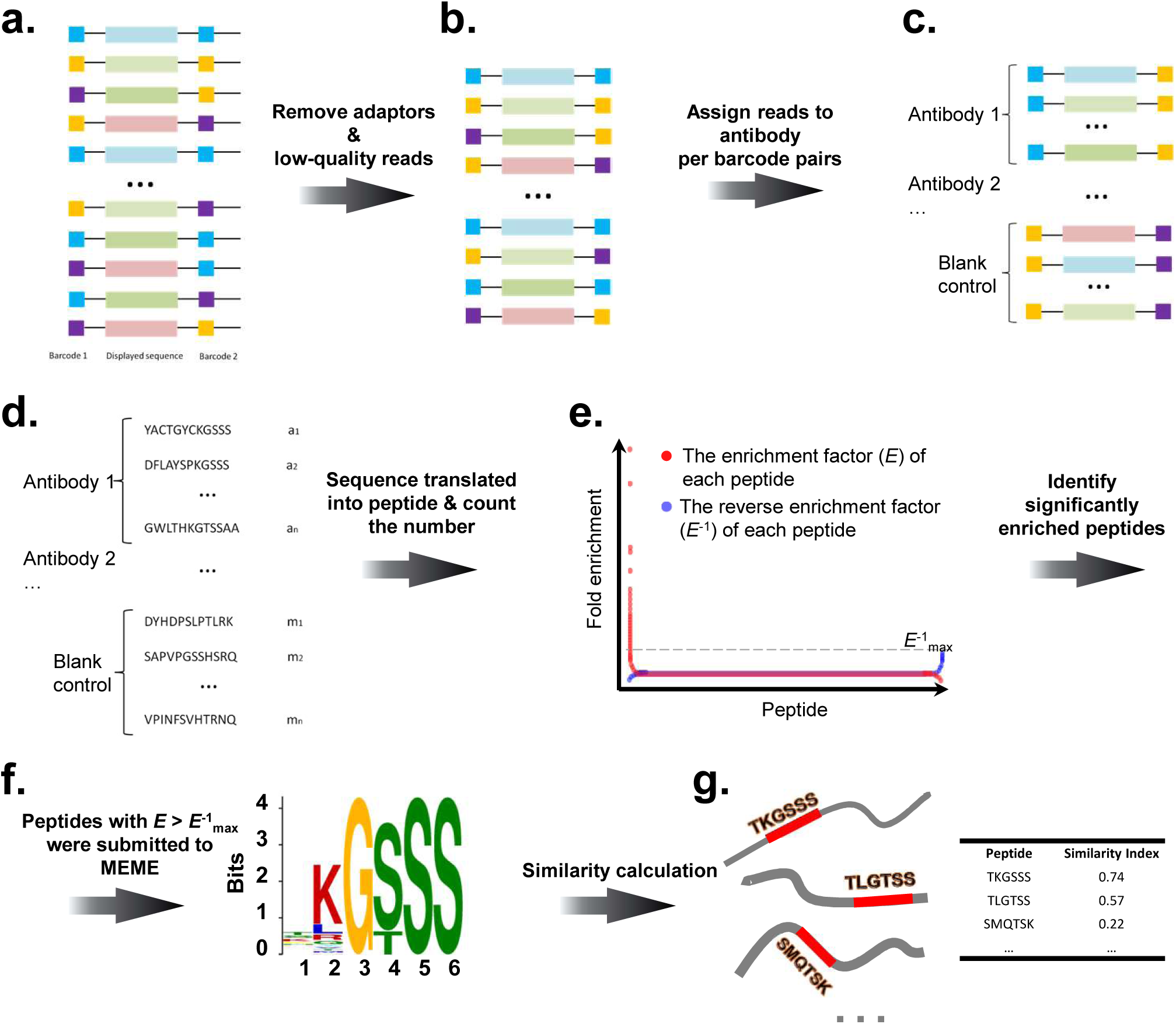
Workflow of data processing. **a**. The structure of the construct for NGS. The solid black lines stand for fixed sequences and colored boxes stand for varied sequences. **b**. The filtered NGS data. **c**. Illustration of reads assigned to antibodies. **d**. Illustration of peptides assigned to antibodies. **e**. The algorithm to obtain enriched peptides. For each peptide, both E (red dots) and E^-1^ (blue dots) are plotted. Only peptides with red dots above the dashed line of the cutoff (E^-1^ max) were kept as enriched peptides for a given antibody. **f**. An example of epitope identified with MEME based on the enriched peptides. The epitope was accompanied by a position-specific probability matrix from MEME. **g**. An example of similarity calculation between the epitopes and the antigens. The red boxes (each consisting of 6 amino acids) in the antigens best matched the position-specific probability matrix (epitope), based on which the similarity index was calculated between the epitope and the antigens.

### Binding peptides/epitopes resolved for widely applied mAbs and pAbs

To test the feasibility of AbMap for monoclonal antibody, we selected two widely applied affinity tag antibodies, *i.e*., anti-6xHis and anti-V5, and went through the procedure depicted in Figure 1 and Figure 2. Figure 3a showed the representative peptides recognized by the anti-6xHis antibody. It is obvious that these peptides are rich of histidine. Because of this, this anti-6xHis antibody was chosen as positive control for the following studies. Clearly, the epitope matched well with 6xHis, however, these six histidines didn’t contribute equally (Figure 3b). A four amino acids epitope was determined for the anti-V5 antibody, which perfectly matches to the V5 sequence (Figure 3b). These results indicate that AbMap is reliable for identifying the epitopes of monoclonal antibodies, and even the epitopes of widely applied tag antibodies could be refined.

**Figure 3.**
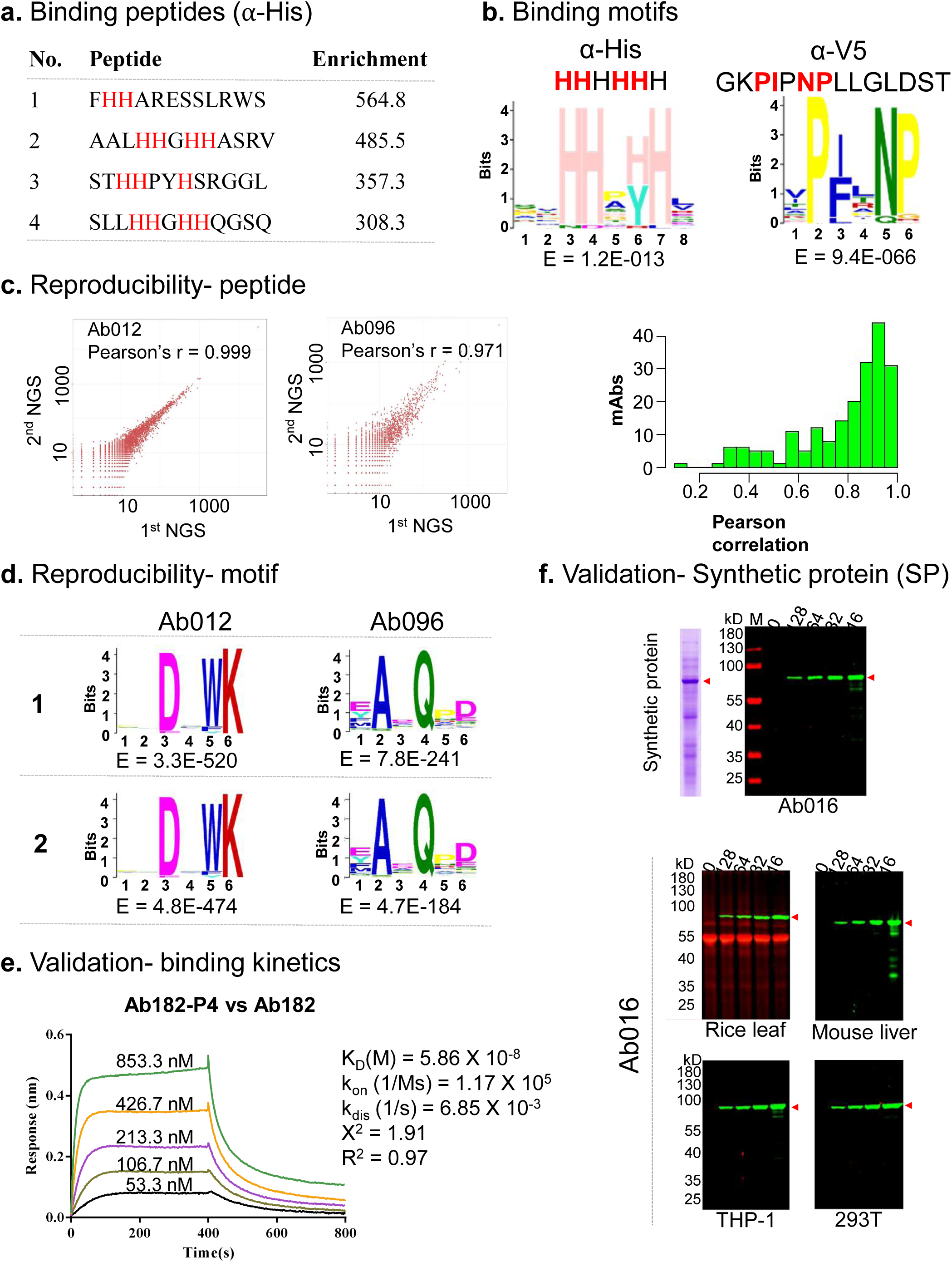
High throughput profiling of antibody binding peptide. **a.** The representative peptides enriched by an anti-6xHis antibody. **b**. The binding epitopes for the anti-6xHis antibody and an anti-V5 antibody. **c**. The reproducibility of AbMap on peptide level. The Pearson correlation of two independent replications of two antibodies (left panel). The summary of the Pearson correlation of the two replications for all the antibodies tested (right panel). **d**. The reproducibility of AbMap on epitope level. Two representative antibodies are shown. **e**. The binding kinetics between representative pair of antibody and its binding peptide. Biolayer interferometry (BLI) was applied for this analysis. **f**. Validation with a *de novo* synthesized protein, *i.e.*, synthetic protein (SP) (see **Figure. S4** for more details). SP was overexpressed in *E. coli*, and the cell lysate was prepared. Western blotting was performed on serially diluted SP in *E. coli* cell lysates. “16” represented 16x dilution. To further assess the specificity of the antibodies, the *E. coli* cell lysate with overexpressed SP was diluted with cell lysates from other species and cells.

Besides monoclonal antibody, polyclonal antibody is also widely applied. To test the possibility of identifying epitope of polyclonal antibodies by AbMap, three antibodies against three well-studied proteins/tags, *i.e*., GST, histone H3 and GFP were chosen. Through the same procedure as shown in Figure 1 and Figure 2, statistically significant epitopes were identified for all these three antibodies. Most of the epitopes matches nicely to the protein sequences. In comparison to monoclonal antibodies, one thing worth noting is that one polyclonal antibody usually recognizes 2-3 different epitopes. These may reflect the polyclonal nature of these antibodies (**Figure S1**).

### High-throughput antibody binding peptide profiling

After the success on several of the widely applied antibodies, we then set to apply AbMap for high-throughput analysis. A total of 202 mouse monoclonal antibodies (**Table S3**) produced from 2012 through immunization of specifically selected peptide antigens were included^21^. The anti-6xHis antibody was set as positive control. Control without the input of antibody was included. Following the procedure as depicted in Figure 1 and Figure 2, we performed the analysis (**Table S4** and **Table S5**). To assess the reproducibility, the binding peptide profiling of the 202 antibodies were repeated twice independently. Pearson analysis showed that between the two repeats, most of the antibodies had a Pearson correlation co-efficient more than 0.8 at the peptide level (Figure 3c and **Table S6**). And almost the identical epitope could be calculated between the two repeats for most of the antibodies (Figure 3d).

It is possible that the antibody enriched peptides are biased to the abundance of the peptide-corresponding phages in the original phage display library. To rule out this possibility, we randomly selected 3 antibodies, and chose 3 peptides that representing high, medium and low enrichment factor for each antibody. The abundance of all these peptides in the original phage library were then quantified by qPCR. The results clearly showed that there was no statistical difference of the abundances among the 3 peptide-corresponding phages for each antibody (**Figure S2**).

It is critical to keep the balance between the NGS reads and the cost. We analyzed the 202 mAbs twice independently, and NGS reads for these two analyses were obtained, *i.e*., 143 M and 75 M. Twelve subsets of reads (from 20 M to 130 M) were randomly selected from the dataset of 143 M reads. Epitopes for the same antibody were compared among all the 14 datasets. The results showed that more than 60 M, *i.e.*, more than 0.3 M reads per antibody was sufficient to assure the successful calculation of the epitopes (**Figure S3**).

To confirm the interaction between antibodies and the enriched peptides, we chose several peptides with varied enrichment factors. The peptides (**Table S7**) were synthesized, biotinylated, and immobilized on a Streptavidin coated sensor. Bio-Layer Interferometry (BLI) assays were performed. The K_D_ could reach to nM or even higher (Figure 3e and **Table S7**).

To further validate more antibody-peptide binding, we selected 43 antibodies, and chose one peptide with enrichment factor about 500 for each of these antibodies (**Table S8**). We linked the sequences of all these peptides together and constructed a synthetic protein (SP), and overexpressed SP in *E. coli* (Figure 3f and **Figure S4a**). To test the sensitivity, SP was serially diluted (**Figure S4b**) and western blotted with all the 43 antibodies individually. The results showed that SP could be clearly and specific recognized by all the 43 antibodies. Even after 128 times dilution, the specific bands were still obviously visible for some of the antibodies (Figure 3f and **Figure S4c**). To further test the specificity, complex samples were prepared by diluting the *E. coli* lysate carrying the overexpressed SP with the lysates from a variety of species and cell lines. We performed western blotting on these complicated samples, using several randomly picked antibodies from the 43 antibodies. The results showed that the sensitivity and specificity of most of the antibodies were consistent even under these complicated background (Figure 3f).

For the high-throughput experiment, as shown in **Figure S5**, 174 antibodies passed the cutoff, *i.e*, with at least one enriched binding peptide. The failure of the 28 antibodies may due to the deleterious effect during the long storage from 2012. Statistically significant epitopes were obtained for 112/174 antibodies. We found that epitopes of 49 out of the 112 antibodies were consistent with the peptides that they were raised against (**Figure S5a**, **Figure S5b** and **Table S5**).

Taken together, we had established a reliable procedure for revealing the antibody recognizing epitope at a high-throughput manner.

### The antibodies were applied to recognize proteins other than the antigens that they were raised against

Based on 3x Cutoff, the epitopes of higher stringency were also calculated (**Table S5**). Given that the multi-specificity nature of antibody (Figure 4a), the epitopes in **Table S5** were used to perform the matching/ homologous searching in human and *E. coli*, as representatives of eukaryotes and prokaryotes. Many proteins were successfully matched (Figure 4b and **Table S9**). The interactions between matched proteins and the corresponding antibodies were validated by western blotting, *i.e.*, human proteins (Figure 4c) and *E. coli* protein (Figure 4d). These results indicate that antibodies recognized the matched proteins at relatively high sensitivity and specificity.

**Figure 4.**
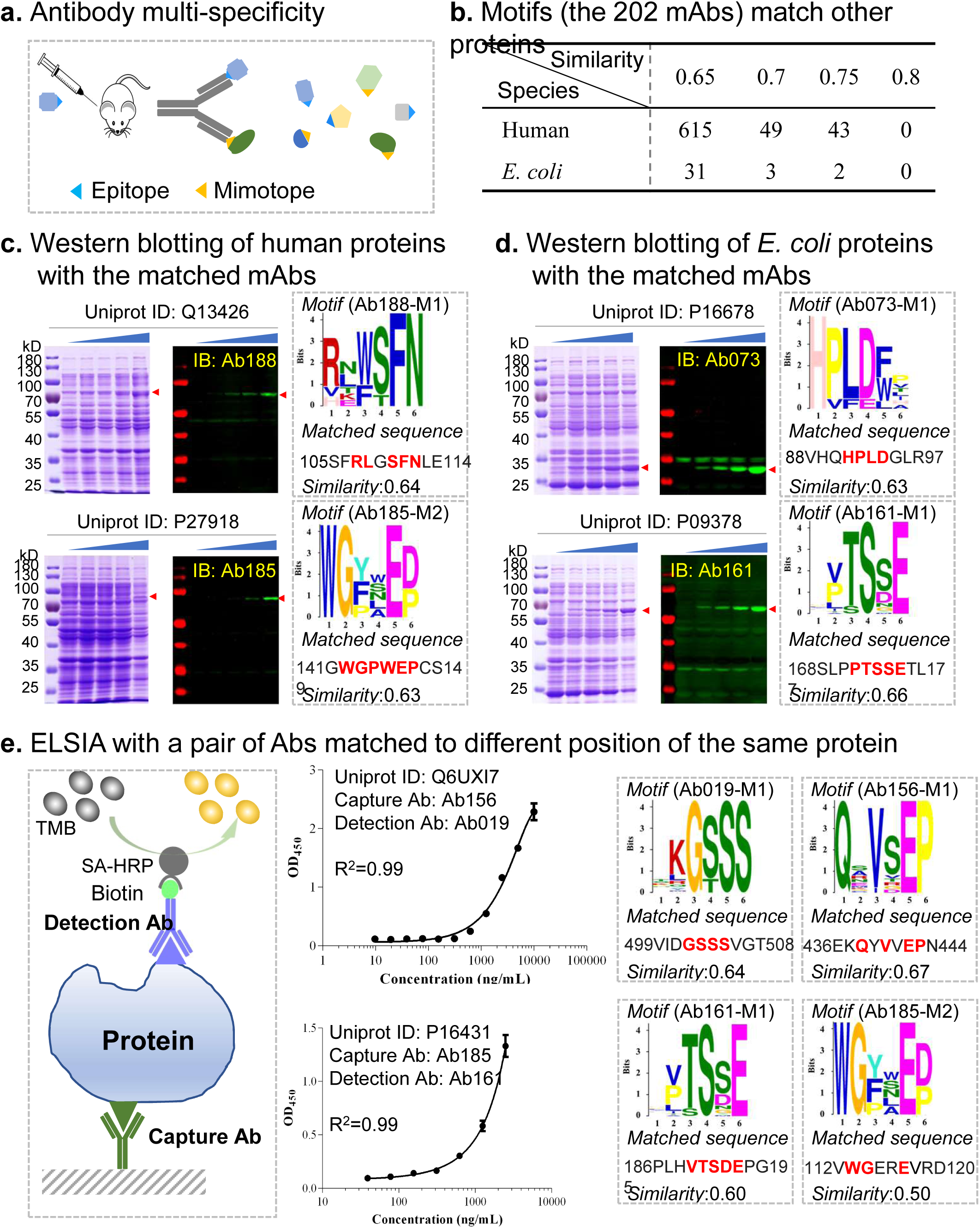
The antibodies could be applied to recognize proteins other than the antigens that they were raised against. **a.** The multi-specific nature of antibody. Usually, an antibody is generated through animal immunization with an antigen of interest. The antibody can recognize specific epitope or mimotope, and this epitope/ mimotope exists on a variety of proteins in nature. **b**. The epitopes revealed through AbMap from the 202 mAbs match to many proteins. **c-d**. Western blotting of proteins with the matched mAbs, human proteins (**c**) and *E. coli* proteins (**d**). **e**. ELISA with a pair of Abs matched to different position of the same protein.

Furthermore, the epitopes of two or more than two antibodies could match to different positions of a protein (**Table S9**), *i.e.*, epitopes Ab019-M1 and Ab156-M1, both are well-matched with human protein Q6UXI7 (Uniprot ID). In another example, epitopes Ab161-M1 and Ab185-M2, are well matched with *E. coli* protein P16431 (Uniprot ID). These antibodies could be paired easily for developing ELISA assays. Relatively high dynamic range and sensitivity were achieved for both two ELISA assays (Figure 4e).

These results demonstrate that guided by the defined epitope, the multi-specificity of an antibody could be explored for potential applications, by targeting the matched proteins other than the antigen that it was raised against.

### Mapping epitopes for anti-PD-1 antibodies

To apply AbMap for clinical related antibodies, we chose anti-PD-1 (human) antibody as the target. Three clinically approved antibodies, *i.e.*, Nivolumab, Pembrolizumab and Sintilimab were included. In addition, we also selected a set of 11 anti-PD-1 antibodies from several producers (Figure 5a). Following the established AbMap procedure, enriched peptides were obtained for all the 14 antibodies, and epitopes were successfully determined for 10 of them (Figure 5b, **Table S10** and **Table S11**). The epitopes of Nivolumab, Sintilimab, ab137132 and 86163T match to PD-1 sequence at high similarity, while the epitopes of the rest of the antibodies match to the PD-1 sequence at relatively low similarity (Figure 5b). The epitope ‘PDR’ of Nivolumab matched perfectly to PD-1, *i.e*., 28-PDR-30, which was within the binding interface between PD-1 and PD-L1 (PDB ID: 5GGR)^19^. To test the effectiveness to predict conformational epitopes based on the linear peptides (**Table S10**), we took advantage of PepSurf^22^ and applied the peptides passed 3x Cutoff as input. Conformational epitopes were predicted for Nivolumab and Pembrolizumab, and these epitopes were highly consistent to the known conformational epitopes of Nivolumab^18^ and Pembrolizumab^19^ (Figure 6). For ab137132 and 86163T, according to the information provided by the producer, the original antigens were protein fragments, not the full-length PD-1. Interestingly, the epitopes of ab137132 and 86163T located exactly in the protein fragments. It was also worth noting that the epitope of ab137132 was very closed to that of Nivolumab and Sintilimab. This suggests that ab137132 could serve as a start point for developing another highly potent therapeutic anti-PD-1 antibody.

**Figure 5.**
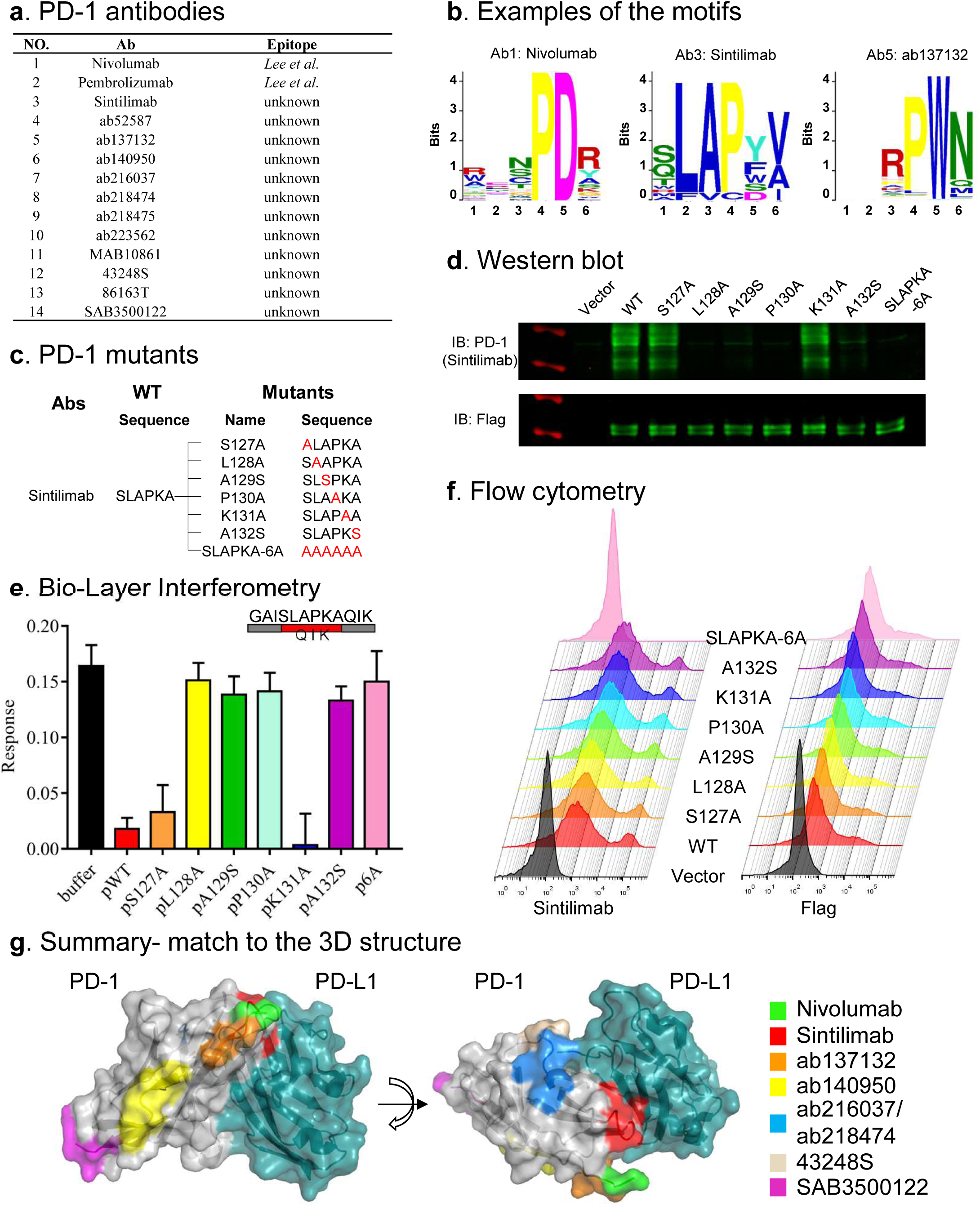
Mapping epitopes for anti-PD-1 antibodies. **a**. The anti-PD-1 antibodies involved in this study. **b**. The resolved epitopes for 3 of the anti-PD-1 antibodies. **c**. A serial of PD-1 mutants were constructed according to the epitope (SLAPKA) of Sintilimab. **d**. Western blotting validation of the epitope of Sintilimab using the PD-1 mutants. **e**. Test the inhibition on the binding between Sintilimab and PD-1 with the WT and mutated peptides by biolayer interferometry (BLI). **f**. Flowcytometry validation of the binding epitope of Sintilimab on PD-1 using the PD-1 mutants. **g**. The distribution of 7 epitopes recognized by 8 anti-PD-1 antibodies on the surface of the co-crystal structure of PD-1 and PD-L1.

**Figure 6.**
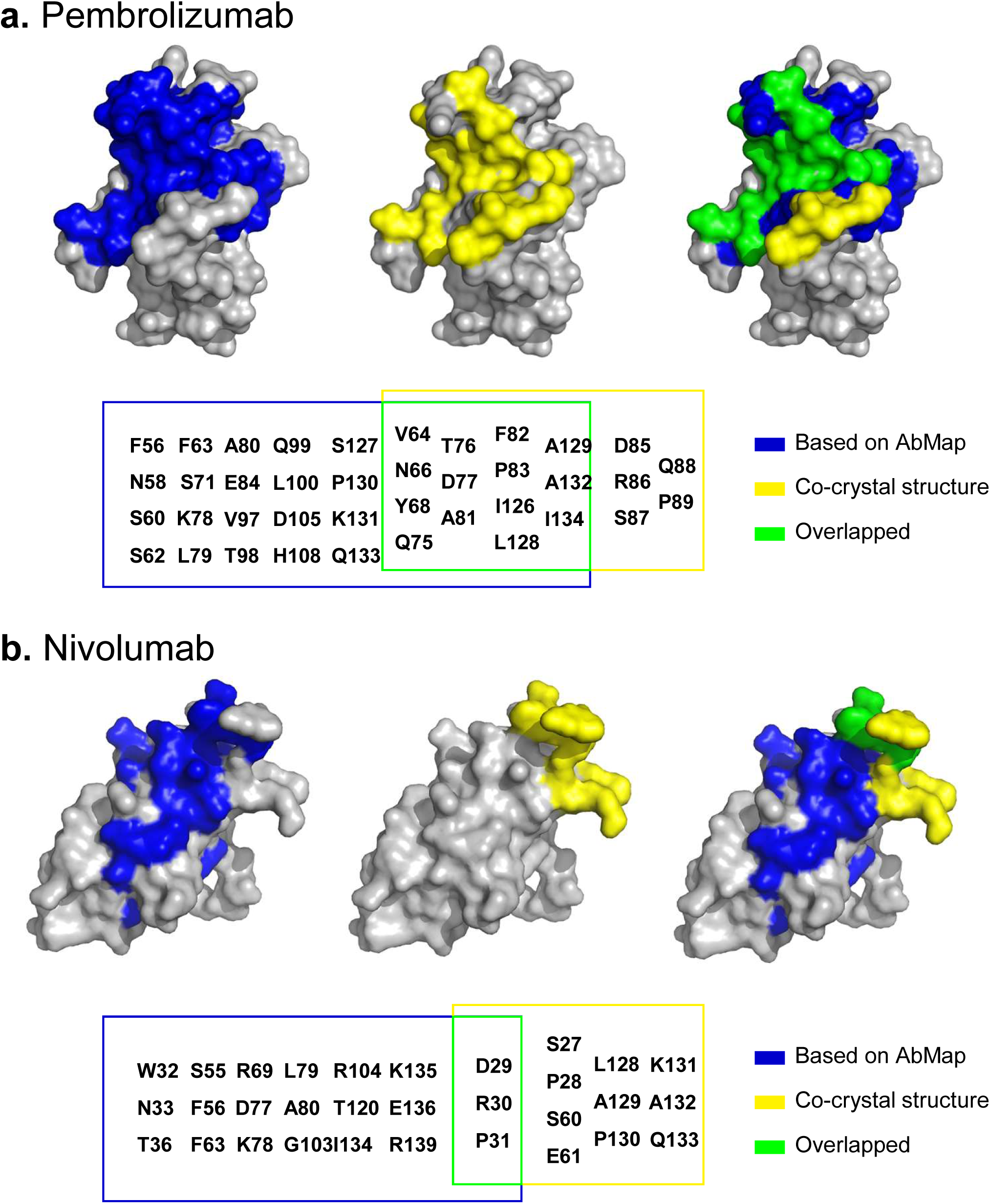
The conformational epitopes predicted based on the AbMap results are consistent to the known antibody-antigen co-crystal structures. **a.** Pembrolizumab **b.** Nivolumab.

Sintilimab had already been approved for therapeutics, however, the binding epitope on PD-1 was still unknown. We decided to use Sintilimab and the epitope as an example for the validation. The epitope (Figure 5b) matched sequence on PD-1 was 127-SLAPKA-132, which also locates at the binding interface of PD-1 and PD-L1. We constructed several mutants of PD-1 by changing the amino acid within 127-SLAPKA-132 (Figure 5c). All the PD-1 mutants were cloned into pFlag-CMV4 and expressed in HEK293T cells. The cell lysates were then denatured and blotted by Sintilimab. The results showed that Sintilimab did not bind L128A, P130A, SLAPKA-6A mutants, while weakly bindings were observed for mutants A129S and A132S mutants (Figure 5d). These results indicate that L128, A129, P130 and A132 were the most critical epitope sites on PD-1 for Sintilimab.

To further validate the epitope of Sintilimab at a conformational setting, BLI assays were performed by incubate the antibody with purified PD-1, upon the addition of the mutated peptides (**Table S7**) to compete the binding. The result was similar to that of WB (Figure 5d). pWT, pS127A and pK131A significantly competed the binding of PD-1 to Sintilimab (Figure 5e).

To confirm the Sintilimab epitope in a more physiological relevant setting, the mutants (Figure 5c) were expressed on the surface of HEK293T cells and detected by flow cytometry (Figure 5f). The results were different from that of WB and BLI. A plausible explanation was that besides L128, A129, P130 and A132, other critical sites yet to be identified may also involve in the binding. Thus, the mutation of only one residue was not significant enough to disrupt the binding between Sintilimab and the native PD-1, while the mutant SLAPKA-6A could (**Figure S6**).

In summary, our results demonstrated AbMap could at least accurately reveal a part, if not all, of the epitope of any antibody at single amino acid resolution. The epitopes of a set of anti-PD-1 antibodies were determined, which were distributed at different locations of PD-1 (Figure 5g, **Movie S1** and **Movie S2**).

## Discussion

We have established the AbMap technology for high-throughput antibody epitope mapping. With AbMap, one technician is able to map more than 200 antibodies in one month at affordable cost. More than 50% of the determined epitopes match to the original antigen. The peptides/epitopes and the corresponding antibodies were explored for a variety of applications, from WB, ELISA and IP. We also map the binding epitopes for 14 anti-PD-1 antibodies, including Nivolumab, Pembrolizumab and Sintilimab. Many of the epitopes match to the sequence of PD-1 either linearly or conformationally, and consistent with the co-crystal structures of antibody and PD-1^18, 19^.

Through AbMap, the profile of the peptides bound by an input antibody could be readily resolved. We argue that the peptide binding profile of an antibody may could represent all the peptides that it could bind in nature. Thus, conceptually, we define an antibody’s binding peptide profile as Binding Capacity (BiC). Based on BiC, we could easily predict what proteins an antibody binds, and precisely on what sites.

There are several advantages of AbMap in comparison to traditional epitope mapping technologies. Firstly, AbMap enables the epitope mapping of hundreds of antibodies simultaneously. For traditional technologies, such as X-ray crystallography^8^, and microarray^12^, usually, only one antibody could be analyzed at a time. By these technologies, it is a common practice to perform epitope mapping only on the antibodies which have the highest potential. In contrast, it is manageable to map 10,000 to even 100,000 antibodies by AbMap. Secondly, the cost of AbMap per antibody is 1-2 order of magnitudes lower than that of traditional technologies. For AbMap, the most expensive part is NGS. In our case, we usually test 200 antibodies in one experiment, and 70 M of NGS reads could assure the reliability. Thus, the NGS cost per antibody is affordable. The low cost enables us, for the first time, to perform on-demand antibody epitope mapping. Thirdly, the accessibility of AbMap is very high. Except NGS, neither complicated technique nor sophisticated instrument is required. The key reagent is the phage library, which is commercially available.

There are also some limitations of AbMap. Firstly, AbMap is intrinsically difficult to map conformational epitopes. The length of the peptides that we used is 12-aa, and it is hard to preserve any effective conformational information. To address this limitation, one solution is to prepare a phage display library with peptides of much longer size, for example 56-aa^23^. Another solution is to predict the conformational epitope through computational tools, such as PepSurf^22^. As demonstrated by the results of Nivolumab and Pembrolizumab, relative correct conformational epitopes could be successfully predicted^18, 19^ (Figure 6). Secondly, the current bioinformatic tools are not optimized for AbMap. It is critical to calculate/predict the linear or conformational epitope after we have the BiC of an antibody. MEME^24^ is used for calculation of linear epitope, and PepSurf is for conformational epitope. For these two tools, the information of peptide abundance is not taken into consideration. To obtain more accurate epitope information from the enriched peptides, specialized bioinformatic tools optimized for AbMap are needed in future studies.

The capability of AbMap for high-throughput and reliable mapping of antibody epitopes at affordable cost makes it an enabling technology. With AbMap, a variety of applications are possible, in the development of both therapeutic antibodies and biological reagents. For therapeutic antibody, we can apply AbMap early in the development pipeline, by comparing the resolved binding epitopes among the antibodies, we can discard the redundant antibodies with similar epitopes and focus limited resource on the unique ones. There is a possibility to produce an antibody recognizes similar epitope as that of the existing therapeutic antibody, thus making the developer to encounter intellectual property infringement^25^. To avoid this, we can compare the epitopes among the candidates and the approved therapeutic antibodies as early as possible. For biological reagents development, AbMap could be applied in two directions: 1. Quality control of the antibodies; 2. Reuse of the existing antibodies. It is possible to resolve the binding peptides/epitopes of any antibody by AbMap, thus we can add another layer of quality control to the antibody. This additional information will definitely guide a researcher to select the right antibody with high confidence. By searching the high confident epitope, an antibody could be re-discovered for testing other proteins. When a large number of antibodies are analyzed, it is highly possible to re-discover antibodies of much better performance than what we showed in Figure 4.

Taken together, facilitated by the high-throughput power of phage displayed peptide library and NGS, we have developed the AbMap technology. AbMap enables, for the first time, the epitopes mapping for hundreds of antibodies simultaneously, at a much less cost than that of traditional technologies, while maintains high accuracy. The identified peptides/epitopes could be explored for a wide range of applications. AbMap technology is highly accessible and generally applicable. We believe that this enabling technology will be applied widely wherever epitope mapping is necessary, and unprecedently accelerate both clinical and basic studies.

## Acknowledgements

This work was partially supported by National Key Research and Development Program of China Grant (No. 2016YFA0500600), National Natural Science Foundation of China (No. 31600672, 31670831, 31370813, 31501054 and 31970130), Open Foundation of Key Laboratory of Systems Biomedicine (No. KLSB2017QN-01). We thank Abmart (Shanghai) for kindly providing the 202 monoclonal antibodies. We thank PTM Biolabs for providing antibodies. We thank Dr. Ai-wu Zhou for providing the recombinant PD-1. We thank Dr. Lan Wang and Dr. Xiaojuan Yu for providing key reagents.

## Author contributions

S.-c. T., X.-d. Z., H. L. and H. Q. designed the study; H. Q. performed the experiments related to the 202 mouse monoclonal antibody; M.-l. M. performed the analysis of PD-1; C. -s. H. and H. L. performed NGS and the data analysis; All the authors analyzed the data; S.-c. T, H. Q., M.-l. M., C.-s. H., X.-d. Z., and H. L. wrote the manuscript.

## Declaration of conflict of interest

The authors declare no conflict of interest.

## Materials and methods

### Enrichment of phage displayed peptides through the binding of antibodies Screening the antibodies’ binding peptides

Briefly, each well of a 96-well PCR plate was blocked with 200 μL of 3% bovine serum albumin in TBST (containing 0.5% Tween 20 in TBS) overnight on a rotator at 4°C. Antibodies (0.02-0.8 μg) of immunoglobulin G (IgG) and 10 μL amplified Ph.D.-12 library (∼100 fold representation, 10^11^ plaque-forming units for a library of 10^9^ clones) were added to each pre-blocked well. The antibodies and phages in Ph.D.-12 library were incubated overnight on a rotator at 4°C. A 10 μL Dynabeads^®^ protein G (Thermo Scientific, CA, USA) were added into each well and to capture the antibody-phage complex for 4 hrs on a rotator at 4°C. With a 96-well magnetic stand, the Dynabeads^®^ protein G were washed three times with 200 μL of TBST and an additional wash with ddw. The beads in each well were resuspended in 15 μL ddw and the phages were lysed at 95°C for 10 min. The phage lysates were stored at −80℃.

### Constructing library for NGS

The library for NGS was constructed following a protocol with slight modifications^23^. Briefly, two rounds of extension PCR were performed by using Q5® Hot Start High-Fidelity DNA Polymerase from New England Biolabs (MA, USA). In the first round of extension PCR, the phage lysate from each well was used as template. The primers used were shown in **Table S1**. A unique combination of up primer (X-SXX-23R) and down primer (X-NXX-18) was set for each antibody. A total of 25 cycles were performed. After evaluated by electrophoresis, the products of first round extension PCR were purified by gel extraction kit (TIANGEN Biotech, Beijing, China), of which 5 μL of each sample was used as the template in the second extension PCR. In the second extension PCR, the primer combination S502 and N701 was used. Besides that, only 10 cycles were performed. The PCR products were evaluated by electrophoresis and purified as that of the first round of PCR. The concentration of each sample was determined by NanoDrop 2000c Spectrophotometer (Thermo Scientific, DE, USA). All the PCR products were mixed with equimolar amounts. The quality of NGS library was checked by TA clone and Sanger sequencing. After examining the structure of each insert and frequency of different barcode pair, the NGS library was subjected to Illumina sequencing with paired-end 2×150 as the sequencing mode.

### NGS data processing and analysis

#### Quality control of NGS data

To filter out the low-quality reads from the raw data (Figure 2a), the software Trimmomatic (version 0.35) was used with the parameter MINLEN set to 150 and other parameters set to default^26^. Since the insert size of our library was exactly 103 bp, the rest part (47 bp) of each read was trimmed as 3’ adaptors (Figure 2b). To ensure high sequencing fidelity, only those paired reads with exact same insert sequence (*i.e.*, no mismatch in the 103-bp region between the paired reads) were kept as clean data for downstream analysis.

### Peptide count normalization

Within each read, there was two 8-bp barcodes and a 36-bp varied DNA sequence. Each pair of 8-bp barcodes uniquely defined an antibody sample where the insert sequence came from (Figure 2c). The 36-bp DNA sequence, which corresponded to the peptide sequence displayed by phage, was translated into a 12-aa peptide and assigned to the corresponding antibody according to the unique barcode pair (Figure 2d). After that, the number of each peptide for each antibody was counted and put into a N x M matrix (defined as raw count matrix) whose rows represented N different 12-aa peptides and columns represented M different antibodies. Thus, each cell of the matrix was the raw count of a peptide. Then the raw count was normalized by the total count of peptides in the column. Additionally, the arithmetic mean count of each peptide was calculated for all control samples, which was further added into the matrix as the control column. The resulting matrix was defined as normalized count matrix.

### Identify enriched peptides for antibodies

Based on the normalized count matrix, the enriched peptides for each antibody were determined by comparison between the antibody column and the control column. Since there was no antibody added into the control samples, all peptide counts observed in the control samples were considered as background noise. For any antibody in the matrix, the enrichment factors (*E*) of all peptides were defined as the antibody column divided by the control column. Due to the background noise, the values in the control column were often much greater than those in the antibody column. Thus, for the same antibody, the reverse enrichment factors (*E*^-1^) were calculated using the control column divided by the antibody column and the maximum reverse enrichment factor (*E*^-1^_max_) was used as the cutoff for the enrichment factors. Unless otherwise specified, only those peptides with *E* > *E*^-1^_max_ were kept as enriched ones for downstream analysis (Figure 2e).

### Epitope analysis for each antibody

Enriched peptides for each antibody were submitted to the standalone version of MEME for epitope discovery using parameters -evt 0.001 -nepitopes 8 -minw 6 -maxw 10 with the rest set to default^24^. Epitopes with E-values <0.05 were considered to be reliable ones for each antibody (Figure 2f).

### Reproducibility analysis

The reproducibility of our data was probed on peptide level. The Pearson correlation coefficient was calculated using the raw peptide counts from two experimental replicates of the same antibody.

### Sequencing depth for epitope discovery

To analyze the impact of sequencing depth on epitope discovery, we performed down-sampling to randomly draw reads from the raw sequencing data (*i.e.*, FASTQ files). We randomly sampled the raw data (143 M paired reads) into 12 data files consisting of 20, 30, 40, 50, 60, 70, 80, 90, 100, 110, 120 and 130 M reads each, performed the same analysis to obtain the epitopes for each antibody, and compare the epitopes with those from the raw data. Based on the epitope reproducibility, the suggested cost-effective sequencing depth was determined for epitope discovery.

### Statistical analyses

All statistical analyses were performed based on the R language (http://www.r-project.org/). In the case of multiple hypothesis testing, *p*-values were corrected with BH method unless otherwise specified^27^.

### Validation the bindings between the antibodies and the peptides

#### Measuring the affinities between antibodies and their binding peptides

All the peptides (**Table S7**) were synthesized by GL Biochem (Shanghai) Ltd (Shanghai, China). According to the manufacturer’s guidelines, the peptides were labelled with biotin by using Sulfo-NHS-LC-Biotin (Thermo Scientific, IL, USA). The unreacted Sulfo-NHS-LC-Biotin was quenched by 1 M glycine solution. The biotinylated peptides were diluted to 2 μM and loaded on pre-wet SA sensor from Pall ForteBio (CA, USA). A serial of concentrations of antibodies were tested against the sensors loaded with peptides. The results were recorded by Octet Red 96 system and analyzed by Octet Software v7.x from Pall ForteBio.

### Validation peptides binding with corresponding antibodies by western blot

A total of 43 antibodies, and one binding peptide for each antibody were selected (**Table S8**). The sequences of these peptides as well as several widely applied affinity tags were linked together, and a synthetic protein (SP) was constructed. The protein sequence of SP was translated into DNA sequence, codon optimized and synthesized by GenScript (Nanjing, China) and integrated into expression vector pET-28a. SP was induced at 0.5 mM IPTG and 16℃ for overnight. A blank control, the *E. coli* containing empty pET-28a, was cultured and induced at the same condition. To prepare the samples, the lysate of *E. coli* containing SP was diluted by two-fold serial dilutions with the lysate of blank control. All the samples were mixed with 5 x loading buffer and boiled for 5 min. Subsequently, the SDS-PAGE and WB were carried out on 10% SDS-PAGE gel for investigating the binding between SP and the 43 antibodies. In WB, the dilution factor of the 43 antibodies were set as 5,000. In addition, the *E. coli* lysates, containing different amount of SP, were mixed with lysates of tissues from a variety of species and lysates of a variety of cell lines for investigating the antibody specificity.

### Applications of the determined peptides/epitopes

#### Matching the epitopes to proteins in human and E. coli

We designed a home-made sliding-window strategy to calculate an epitope-proteins similarity index as follows. First, we used MEME to generate a position-specific probability matrix that showed the frequency of each amino acid at each position of the identified epitope^24^. Second, we used this matrix as a sliding window (by one amino acid at a time) to scan the peptide from N terminal to C terminal and locate which part of the peptide best matched the matrix. Third, based on the best matched part, we calculate the similarity index for the whole proteins.

The epitopes of the 202 antibodies were matched to immunogen and all the proteins in Human and *E. coli*. A list of proteins from Human and *E. coli* were revealed, with high sequence similarity to the identified epitopes.

### Validating the binding between antibodies and proteins by western blotting

Proteins with similarities bigger than 0.5 with epitopes, were chosen for further validation. In the human ORF library from Invitrogen, the plasmids carry the genes of the selected proteins were transferred to the expression vector pDEST^TM^ 15 by LR reactions (Thermo Scientific, CA, USA). Subsequently, all the reactions were transformed to BL21 (DE3) chemically competent cell and confirmed by Sanger sequencing. All the proteins were induced with 0.5 mM IPTG at 37℃ for 2 hrs. All the samples were used for western blotting for validating the binding between the antibodies and the target proteins. The dilution factor of antibodies was 1,000 in western blotting.

### Antibody paired for ELISA

#### Proteins purification

The gene for protein Q6UXI7 was amplified with the primers F30B and R30B (**Table S2**), which contained the restriction sites of *Eco* RI and *Hind* III, respectively. The amplified fragment and expression vector pET-28a were digested with *Eco* RI and *Hind* III. After purification, the gene fragment and the linearized vector were linked by T4 DNA ligase (New England Biolabs, MA, USA). Subsequently, the plasmid was transformed into BL21 (DE3) chemically competent cell. The final expression vector containing the gene for protein Q6UXI7 was confirmed by Sanger sequencing. Proteins Q6UXI7 and P16431 were induced by 0.5 mM IPTG at 37℃ for 2 hrs. The cells were harvested and lysed by lysis buffer (TBST containing 2% SDS). The proteins were purified on the Ni-NTA affinity columns. The purified proteins were stored at −80℃.

### Biotinylation of the antibodies

Ab019 and Ab161 were purified by incubation with Protein G beads and their concentrations were determined by BCA protein assay kit (Thermo Scientific, IL, USA). According to the manufacturer’s manual, Ab019 and Ab161 were labelled with Sulfo-NHS-LC-Biotin (Thermo Scientific, IL, USA). The free Sulfo-NHS-LC-Biotin were removed by dialysis. The concentrations of the biotinylated antibodies were determined by BCA.

### ELISA

The first antibody was diluted to 4 μg/mL with PBST (PBS containing 1% Triton X-100). The 96-well plates were sensitized with 50 µL antibody per well. After centrifugation for 2 min at 550 g for distributing proteins evenly, the 96-well plates were incubated at 4℃ for overnight. Subsequently, the plates were emptied and blocked with 300 µL blocking buffer (Thermo Scientific, IL, USA) at RT for 1 hrs. The plates were washed three times by PBST and were incubated with 50 µL target protein at different concentration at RT for 1 h. The plates were further washed with PBST for another three times, and incubated with 50 µL biotinylated antibody at the concentration of 2 μg/mL. After incubation for 1 hrs at RT, the plates were further washed with PBST for three times and incubated with 50 µL 1000X diluted SA-HRP (Sigma-Aldrich, MO, USA). After 3 washed with PBST, each well was added with 50 µL TMB solution (Sigma-Aldrich, MO, USA) and incubated for 20 min at 37℃. The plates were incubated with 50 µL of 1 M H_2_SO_4_ to stop the reaction. The results were recorded with a multichannel spectrophotometer (BioTek Instruments, VA, USA) at 450 nm. Three replicates were carried out. A typical third order polynomial (cubic) nonlinear regression model was used for standard curve-fitting for ELISA, which was further used to calculate the concentration of target protein in samples.

### Applications related antibodies for PD-1

#### Obtaining the epitopes for a set of anti-PD-1 antibodies

Three approved anti-PD-1 antibodies (Nivolumab, Pembrolizumab and Sintilimab) and 11 anti-PD-1 antibodies from several renown antibody companies were selected, including ab52587, ab137132, ab140950, ab216037, ab218474, ab218475, ab223562 (abcam, MA, USA), MAB10861 (R&D systems, MN, USA), 43248S, 86163T (CST, MA, USA), SAB3500122 (Sigma-Aldrich, MO, USA). These antibodies only recognize human PD-1 and were all monoclonal antibodies. All the binding peptides and epitopes of each antibody were obtained by following the procedure of AbMap. Given that the available structure of PD-1, conformational epitopes were predicted by Algorithm PepSurf (http://pepitope.tau.ac.il/). Antibodies binding peptides with enrichment factor pass the 3x Cutoff and PD-1 structure file (PDB ID: 5GGS) were submitted. The gap penalty value was set to −5, and the probability for obtaining the best path was set to Default. The predicted cluster with highest score was further analyzed.

### Constructing the mutants of PD-1 for Sintilimab

According to the epitope for Sintilimab, PD-1 mutants (S127A, L128A, A129S, P130A, K131A, A132S, and SLAPKA-6A) were constructed and introduced in pFlag-CMV4-PD-1 by QuikChange Site-Directed Mutagenesis. All constructs were confirmed by sanger sequencing. All primers were listed in **Table S2**. The thermal profile was followed: denaturing at 95℃ for 2 min, 25 cycles (95℃, 30; 55℃, 30 s; 72℃, 4 min), and further extension at 72℃ for 10 min.

### Inhibition effect of peptides on interaction between PD-1 and Sintilimab by BLI

The peptides were incubated with Sintilimab for 1 hrs at RT. The recombinant PD-1 protein was loaded to the surface of prewet NTA biosensors from ForteBio (Pall ForteBio, CA, USA). Subsequently, the NTA biosensors were used to monitor the inhibition effect of peptides on interaction between PD-1 and Sintilimab. All the data were recorded and processed on the Octet Red 96 system from Pall ForteBio.

### Validating epitopes of by Sintilimab FACS

HEK293T were transfected with PD-1 WT and mutant plasmids (S127A, L128A, A129S, P130A, K131A, A132S, and SLAPKA-6A). Cells were separated with 1 mM EDTA-PBS solution after 48 hrs, and incubated with 10 μg/mL of Sintilimab for 30 mins at 4℃, after 3 washes with PBS, cells were incubated with 5 μg/mL of the FITC-labeled secondary antibody for 30 mins at 4℃ in dark, after 3 washes, the cells were analyzed by flow cytometry using LSR Fortessa (BD BioSciences, CA, USA).

## Supplementary materials

### Supplementary figures

**Figure S1. Multiple epitopes are determined for polyclonal antibodies.**

**Figure S2. The enrichment factor of the peptides is not biased to the abundance of the corresponding phage in the phage library. a.** Three antibodies were randomly selected. Three peptides represent high (H), medium (M) and low (L) enrichment factor for each antibody were chosen. Specific upstream primers were designed for the coding DNA sequences of these peptides. **b.** The design of the qPCR primers. A common downstream primer was designed for all the peptides. **c.** The qPCR results (mean ± SD., n = 3 independent experiments).

**Figure S3. Reads required for reliable determination of epitopes.** The sample for secondary biological repeat for 202 mAbs was sequenced twice, generating 143 M and 75 M effective NGS reads each (red marked, real sequencing datasets). Subsets of reads were randomly selected from the dataset of 143 M, from 20 M to 130 M (shadowed, 12 simulated datasets). Epitopes were calculated for all the 202 mAbs in all the datasets. Epitopes for the same antibody were compared among all the 14 datasets. Five representative antibodies were shown.

**Figure S4. Sensitivity test. Western blotting of the 43 antibodies on SP. a.** The composition of protein SP. b. The procedure of sample preparation by serial dilution of SP. c. Western blotting was performed on serially diluted cell lysate, and “8” represents a dilution factor of 8.

**Figure S5. The consistency of the epitopes and the original immunogens that the antibodies were raised against. a.** An example: the epitope of Ab077 perfectly matched to the sequence of the original immunogen. **b.** The summary of AbMap results for the 202 antibodies.

**Figure S6.** The binding mechanism between Sintilimab and PD-1. **a.** Linear epitope. **b.** Conformational epitope.

## Supplementary Tables

**Table S1. Barcodes, indexes used in this study.**

**Table S2. Primers used in this study.**

**Table S3. Details of the 202 mAbs for high-throughput epitope mapping.**

**Table S4. The peptide binding profiles of the 202 antibodies.**

**Table S5. The resolved epitopes for 202 antibodies.**

**Table S6. Reproducibility of the peptide binding profiles of all the 202 antibodies.**

**Table S7. The peptides synthesized in this study.**

**Table S8. The 43 peptides for constructing SP.**

**Table S9. Match the epitopes to proteins from a variety of species.**

**Table S10. The peptide binding profiles of the anti-PD-1 antibodies.**

**Table S11. The resolved epitopes for the anti-PD-1 antibodies (3x Cutoff).**

**Movie S1. The distribution of 7 epitopes recognized by 8 anti-PD-1 antibodies on the surface of the crystal structure of PD-1.**

**Movie S2. The distribution of 7 epitopes recognized by 8 anti-PD-1 antibodies on the surface of the co-crystal structure of PD-1 and PD-L1.**

