## Supplementary material for "Antibody binding epitope Mapping (AbMap) of two hundred antibodies in a single run": Table S1,S2,S3,S4,S5,S6,S7,S8,S9,S10,S11, Figure S1,S2,S3,S4,S5,S6 and Movie S1,S2: Supplemental figures.pdf

| Antibody | Motifs | Protein sequence |
|---|---|---|
| α-GST (Sigma Cat. No. G7781) | 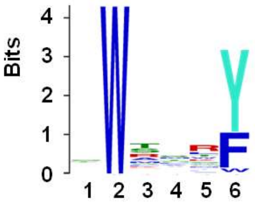 E = 1.2E-55 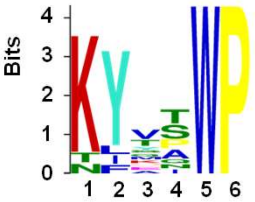 E = 1.6E-6 | **GST protein (Uniprot ID: P08515)**<br>MSPILGYWKIKGLVQPTRLLLEYL<br>EEKYEEHLYERDEGDKWRNKKF<br>ELGLEFPNLPYYIDGDVKLTQSMA<br>IIRYIADKHNMLGGCPKERAEISML<br>EGAVLDIRYGVSRIAYSKDFETLK<br>VDFLSKLPEMLKMFEDRLCHKTY<br>LNGDHVTHPDFMLYDALDVVLYM<br>DPMCLDAFPKLVCFKKRIEAIPQID<br>KYLKSS**KY**IA**WP**LQG**W**QAT**F**GGG<br>DHPPK |
| α-H3 (Abcam Cat. No. ab1791) | 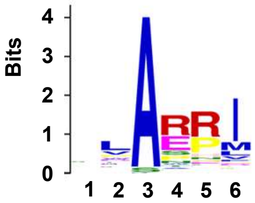 E = 1.1E-62 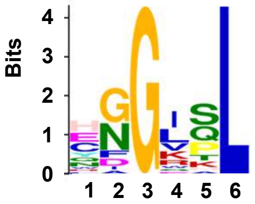 E = 5.5E-10 | **H3 protein (Uniprot ID: P68431)**<br>MARTKQTARKSTGGKAPRKQLAT<br>KAARKSAPATGGVKKPHRYRPGT<br>VALREIRRYQKSTELLIRKLPFQRL<br>VREIAQDFKTDLRFQSSAVMALQ<br>EACEAYLVGLFEDTNLCAIHAKRV<br>TIMPKDIQ**LARRI**RGERA<br><br>**Patial KLH (Uniprot ID: Q6KC56)**<br>…701AMPNKIYDYENVLHYTYED<br>LTF**GGISL**ENIEKMIHENQQEDRIY<br>AGFLLA750… |
| α-GFP (Abcam Cat. No. ab290) | 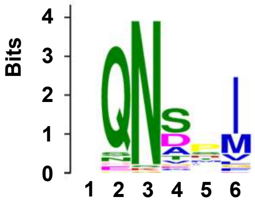 E = 4.0E-39 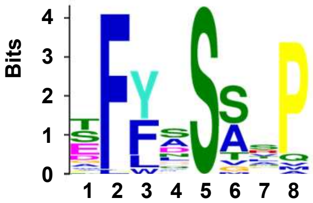 E = 9.4E-35 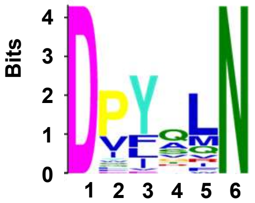 E = 5.2E-35 | **GFP protein (Uniprot ID: P42212)**<br>MSKGEELFTGVVPILVELDGDVNG<br>HKFSVSGEGEGDATYGKLTLKFICT<br>TGKLPVPWPTLVTTFSYGVQCFSR<br>YPDHMKQHD**FF**K**SA**M**P**EGYVQER<br>TIFFKDDGNYKTRAEVKFEGDTLVN<br>RIELKGIDFKEDGNILGHKLEYNYN<br>SHNVYIMADKQKNGIKVNFKIRHNI<br>EDGSVQLA**D**H**YQQN**TP**I**GDGPVLL<br>PDNHYLSTQSALSKDPNEKRDHMV<br>LLEFVTAAGITHGMDELYK |

**Qi *et al.*, Figure S1**. Multiple motifs are determined for polyclonal antibodies.

**a.**

| Antibody | ID of peptide | Sequence of peptide | Enrichment factor | Upstream primer | Primer sequence (5'→3') |
|---|---|---|---|---|---|
| Ab069 | Ab069-P1 | TAQANYVAALQS | 6488.56 | Ab069-H | GAATTATGTGGCGGCTTTGCAG |
| | Ab069-P2 | NAIVSDYYDVLT | 898.35 | Ab069-M | TTCGGATTATTATGATGTGCTGACG |
| | Ab069-P85 | KVTPYMNYVAAL | 58.93 | Ab069-L | TTATATGAATTATGTGGCTGCGTTG |
| Ab099 | Ab099-P1 | SMQLSAVPSSSR | 6403.32 | Ab099-H | GTCTGCGGTGCCTAGTAGTAG |
| | Ab099-P2 | TSSVQMGSVPTY | 3738.01 | Ab099-M | GCAGATGGGTTCGGTGCC |
| | Ab099-P528 | WYPGRQSYAVPP | 73.98 | Ab099-L | TAGGCAGAGTTATGCTGTTCCGC |
| Ab139 | Ab139-P1 | IWARVSEFTRPF | 4391.37 | Ab139-H | GGTGTCTGAGTTTACGAGGCC |
| | Ab139-P2 | GILDRQPGNRPW | 1293.70 | Ab139-M | TCGTCAGCCTGGTAATCGTCC |
| | Ab139-P210 | SAKDAVLITRAW | 123.30 | Ab139-L | TGCTGTGCTTATTACGAGGGCTTG |

**b.**

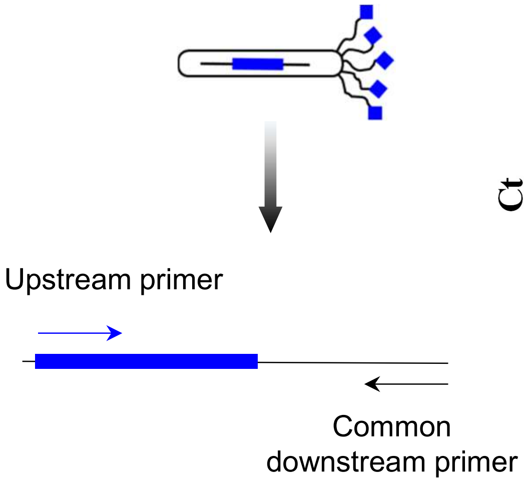


Common downstream primer sequence:
5'-CAGCCCTCATAGTTAGCGTAACG-3'

**c.**

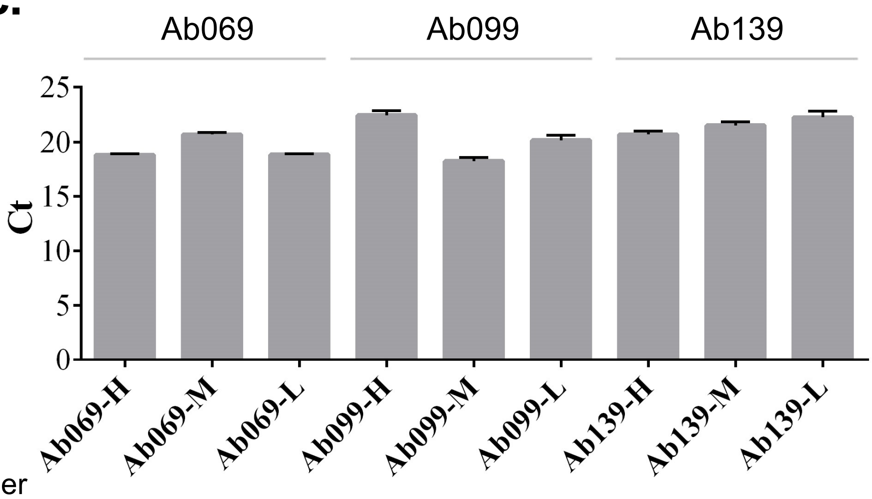


**Qi *et al.*, Figure S2**. The enrichment of the peptides is not biased to the abundance of the corresponding phage in the phage library.

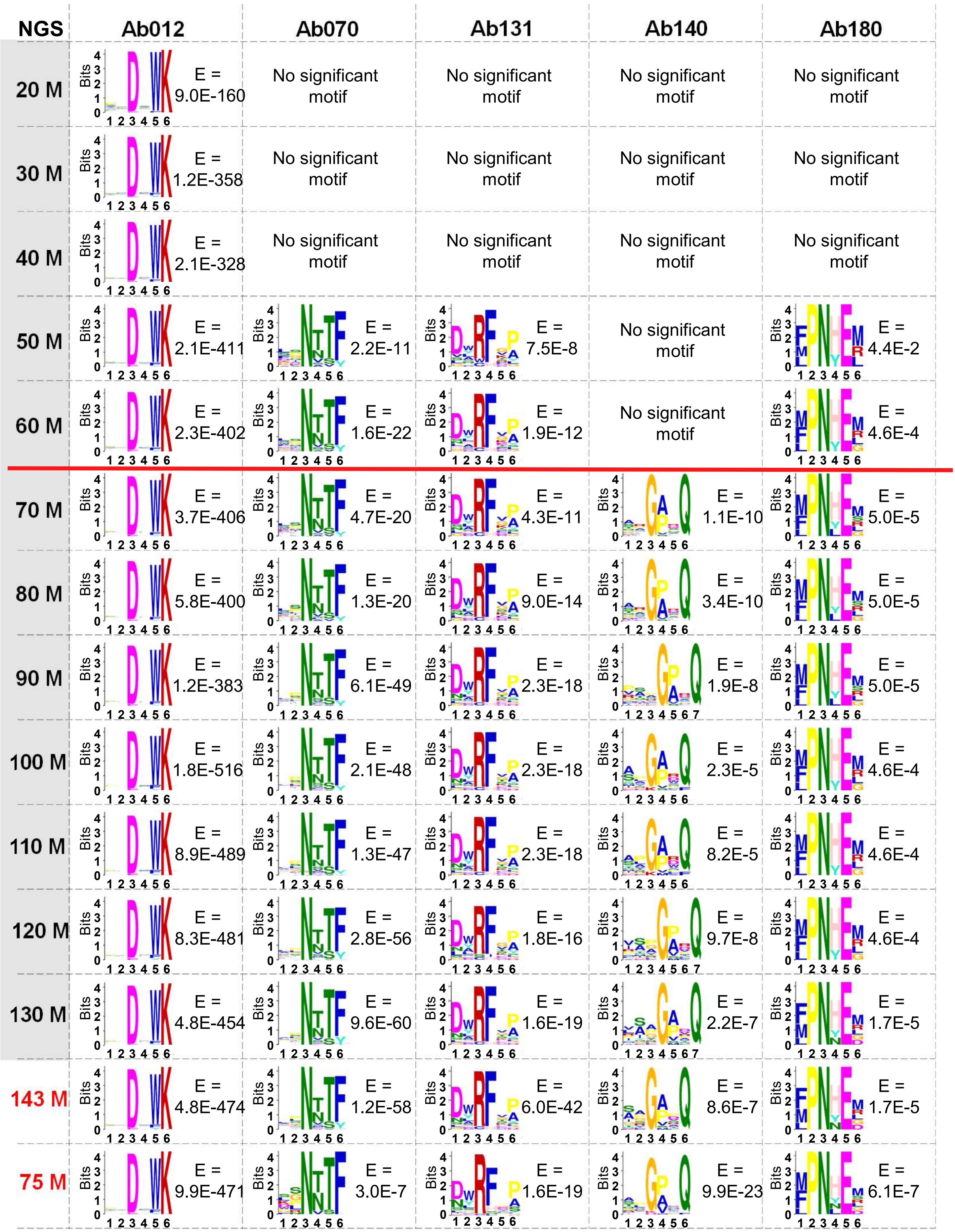


**Qi *et al.*, Figure S3**. Reads required for reliable determination of motifs.

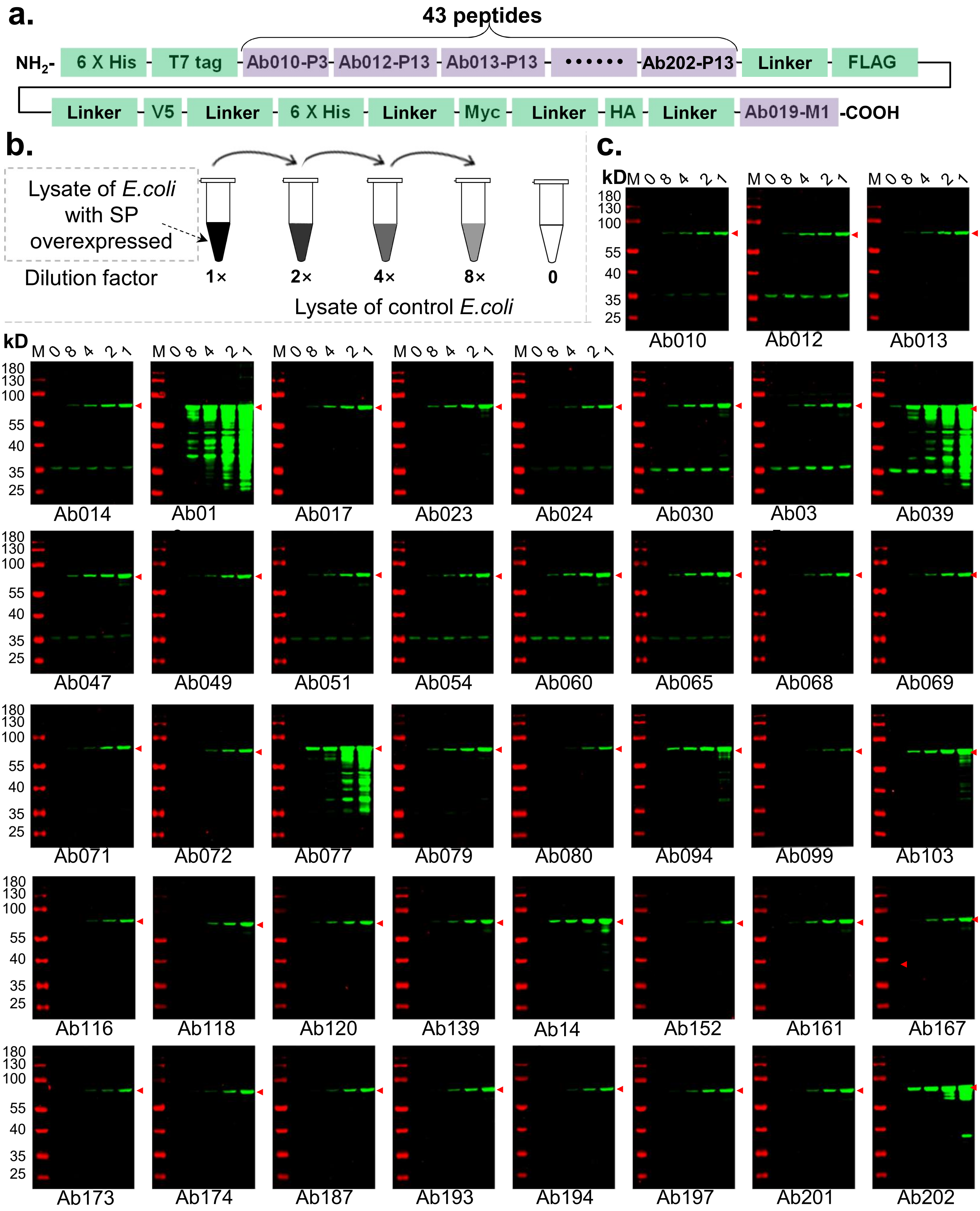


**Qi *et al*., Figure S4**. Sensitivity test. Western blotting of the 43 antibodies on SP.

**a.**

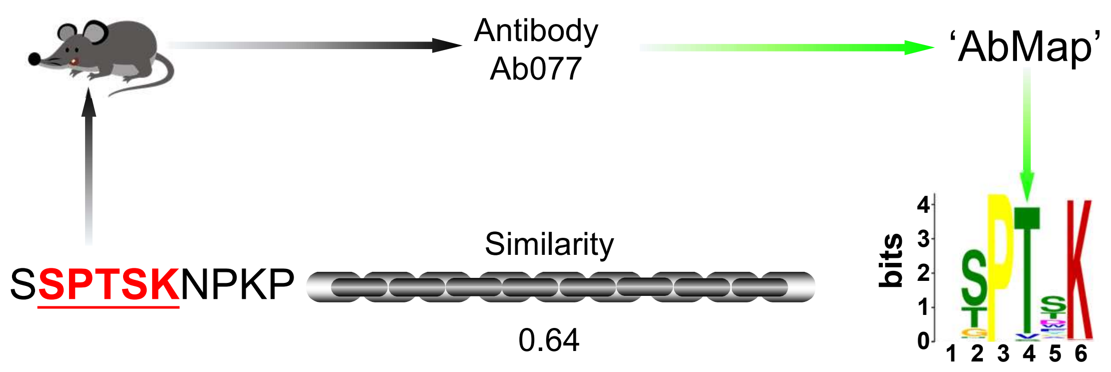


**b.**

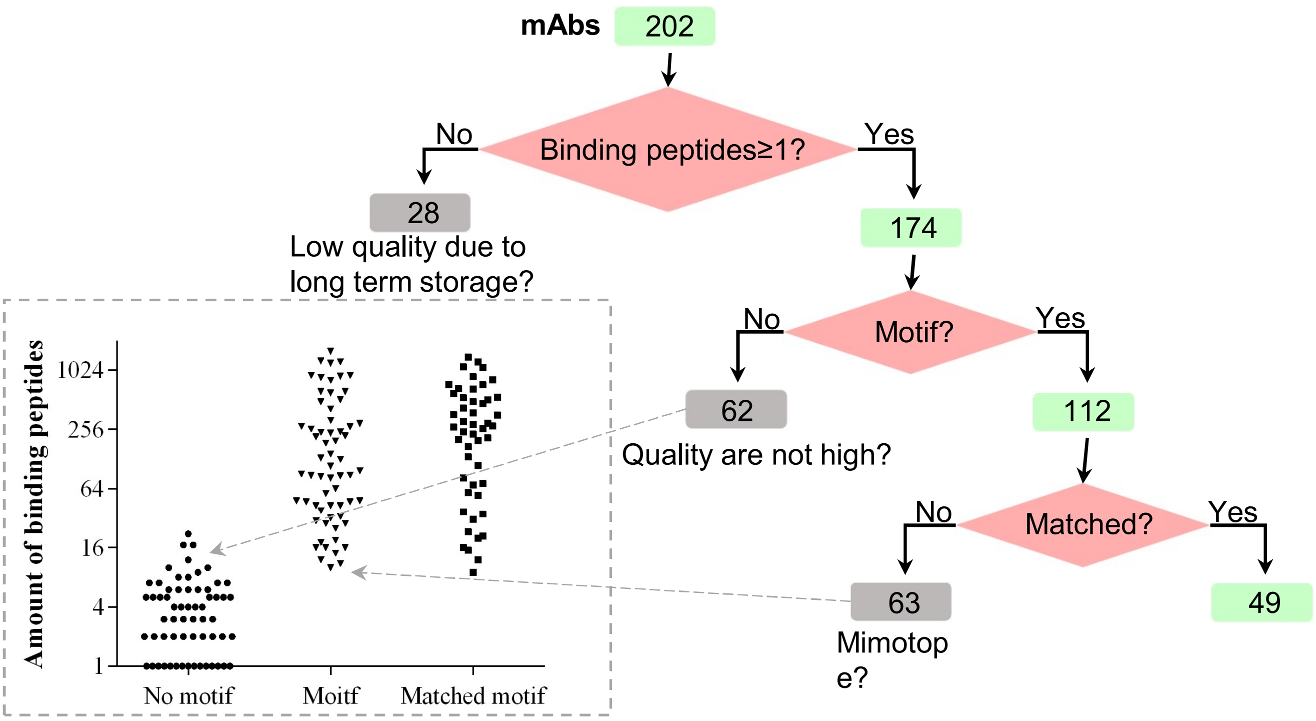


**Qi *et al*., Figure S5**. The consistency of the motifs and the original immunogens that the antibodies were raised against

**a.** Linear epitope. Sintilimab interacts with PD-1.

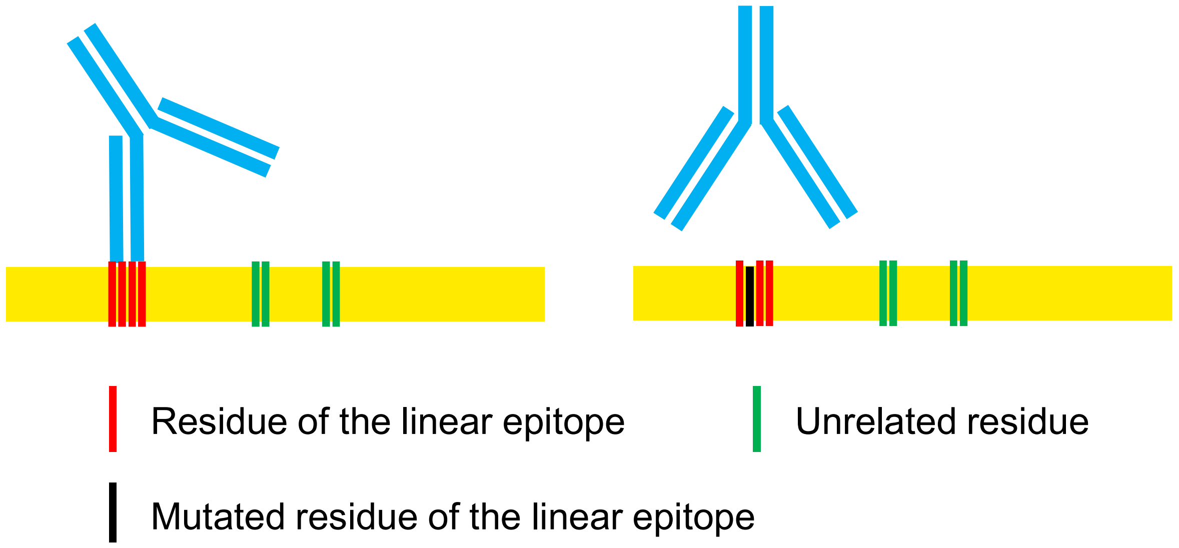


**b.** Conformational epitope. Sintilimab interacts with PD-1.

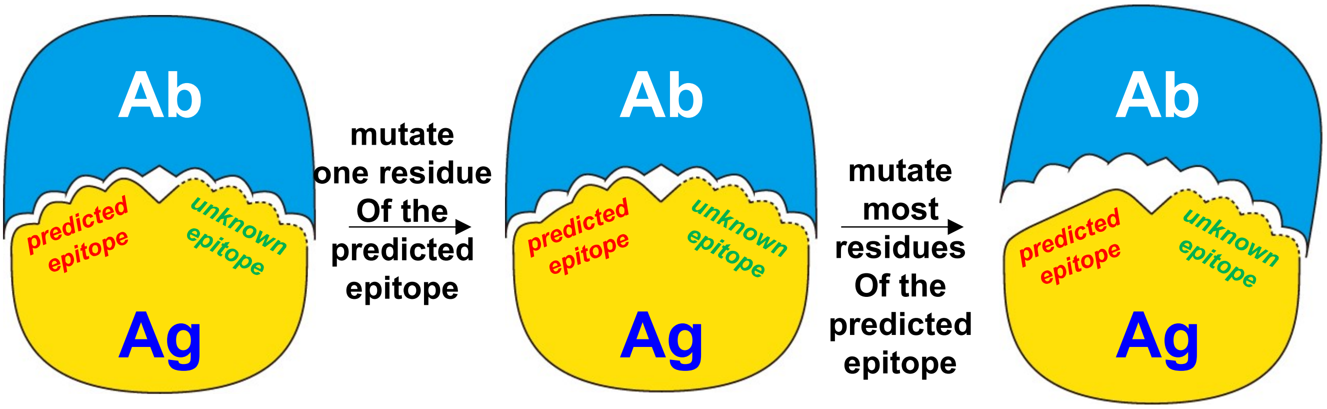


**Qi *et al*., Figure S6.** The binding mechanism between Sintilimab and PD-1.
